# Predicting single mutation effects on binding affinity at protein-protein interfaces based on MMPBSA calculations

**DOI:** 10.64898/2026.09.22.753561

**Authors:** Jakob Noske, Thibaud Lepoivre, Maxim Janzen, Birte Höcker

## Abstract

Accurate prediction of mutation-induced changes in protein-protein affinity remains a central challenge in computational biophysics and protein engineering. Here we present a physics-based scoring method for predicting the effects of single amino acid substitutions on protein-protein and protein-peptide binding affinity. The method is based on MMPBSA energy calculations and implemented in a user-friendly, publicly available workflow (https://github.com/Hoecker-Lab/mmpbsa_scoring). To derive the scoring function, MMPBSA energy terms were fitted using a large and diverse dataset of experimentally measured binding affinity changes upon point mutation. The terms were then evaluated on independent systems. Specifically, we assessed the ability to predict mutational effects at the interface of SARS-CoV-2 spike protein with its cellular receptor, as well as computationally predicted Armadillo-repeat protein-peptide complexes. The approach performs well across diverse targets and shows promising predictive performance on both experimentally determined and computationally predicted structures. It matches or exceeds the accuracy of established physics-based protein design software and state-of-the-art deep learning-based scoring functions when applied to data outside their training domain. The method provides a computationally efficient and interpretable framework for prioritizing interface mutations in protein engineering and binding affinity optimization.

## Introduction

Single residue mutations at protein-protein interfaces can significantly alter the affinity between two binding partners. Predicting this effect remains a fundamental challenge in computational protein design and molecular engineering. Although single residue mutations often affect only a small region of an interface, even subtle changes in molecular interaction networks or conformational dynamics can produce substantial changes in binding free energy (ΔΔ*G_Bindin_*_g_)^1,2^. While protein design efforts have targeted large interfaces achieving affinity via a significant number of newly introduced interactions^3–6^, analyzing the energetics of individual amino acid mutations at binding interfaces can provide important insights into the accuracy of scoring functions and help guide their development^7–9^. Accurate prediction therefore requires both realistic structural models of mutant complexes that reflect the conformational changes and a scoring function capable of resolving small energetic differences.

A wide range of approaches have been developed to address this problem including empirical force fields, knowledge-based potentials, but also machine learning-based scoring functions^9–17^. While recent deep learning models have achieved impressive predictive performance on their respective training sets, they often generalize poorly. Limited availability and heterogeneous quality of experimentally determined datasets can further complicate both model development and benchmarking^18–21^. As a result, there remains considerable interest in interpretable, physics-based energy terms that can be applied across diverse protein systems.

Physics-based scoring functions implemented in protein design methods such as Rosetta^22^, Osprey^23^ or PocketOptimizer^24^ enable efficient evaluation of large numbers of mutations, but often lack accuracy on the single residue level^7^. Their simplified energy terms may not fully capture mutation-dependent changes in electrostatic and solvation effects^7,10,23,25^. In contrast, Molecular Mechanics Poisson-Boltzmann Surface Area (MMPBSA) calculations provide a conceptually simple, physically founded framework that explicitly accounts for these contributions. MMPBSA methods have been successfully applied to estimate binding free energies ^26–29^, and offer an attractive basis for predicting residue-level affinity contributions^9,30^.

Here, we present a MMPBSA-derived scoring function for predicting mutation-induced changes in protein-protein binding affinity. The model combines MMPBSA energy components and folding stability terms through linear regression fitted on a filtered subset of the SKEMPI v2.0 database^31^. To evaluate its robustness and transferability, we tested the resulting scoring function on two independent datasets, namely the Receptor Binding Domain (RBD) of the SARS-CoV-2 spike glycoprotein in complex with the human ACE2 receptor^32^ and a series of designed Armadillo Repeat Proteins (dArmRPs), which offer a modular peptide-binding interface^7,33^. Performance was compared with established physics-based and deep-learning approaches to assess the accuracy and generalizability of the proposed method.

## Results and discussion

The SKEMPI v2.0 database consists of a large set of mutational effects on protein-protein binding affinity. We used a filtered subset of this database to fit our scoring function (see methods). This subset includes ΔΔ*G_Bindin_*_g_ data for a wide variety of diverse single amino acid substitutions at various types of protein-protein interfaces, which should enable the generation of a generalizable and widely applicable scoring function. We calculated the MMPBSA-based energy components and additionally included the folding stability change as calculated by FoldX^10^ (see methods). A least squares linear regression model was then fitted to determine the contribution of each energy term (*w*_1_ to *w*_4_) to the total binding free energy change upon mutation. In the fitted model *w*_0_ represents the intercept of the fit. Relying on a simplified linear regression model instead of a more complex machine learning-based architecture has the advantage that it allows for a direct interpretation of the different energy contributions in the final fitted model. Furthermore, if an energy component is not well-defined in general or not correctly reflected in our model it allows for a more straightforward cancellation of errors as it appears for both the mutant and the wildtype in Δ*G_Bindin_*_g_.

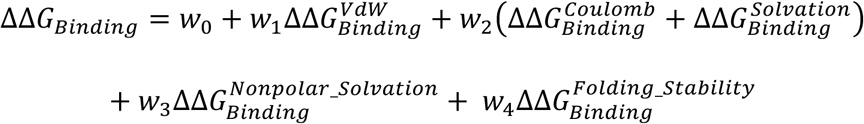

The weights of the fitted model are given in table 1. Both the Van-der-Waals (VdW) and the electrostatic (Coulomb + Solvation) binding free energy component make a significant contribution to the model. The nonpolar solvation energy only makes a very minor contribution, which is in agreement with previous MMPBSA analyses on mutational effects at protein-protein interfaces^11^. The folding stability change significantly contributes to the model as well, which highlights that binding effects of mutations are also heavily influenced by stability changes caused to the protein monomers. This term is typically not considered in MMPBSA calculations as the intramolecular energies normally cancel themselves out. However, our analysis indicates that it is highly relevant and should be considered when calculating ΔΔ*G_Bindin_*_g_ at protein-protein interfaces.

**Table 1:** Coefficients of weights of the fitted linear regression model.

| Weight | Fitted coefficient |
| --- | --- |
| $w_0$ (Intercept) | 0.495 |
| $w_1$ (VdW) | 0.206 |
| $w_2$ (Coulomb + Solvation) | 0.254 |
| $w_3$ (Nonpolar Solvation) | 0.022 |
| $w_4$ (Folding Stability) | 0.277 |

The results obtained after fitting our model (MMPBSA-based) on the SKEMPI v2.0 dataset are shown in figure 1A. To compare our model with an existing physics-based protein design software we additionally evaluated all point mutation effects using the most recent version of the FoldX molecular modelling suite, which includes the recently published FoldX 5.1 force field^10^ (figure 1B). Furthermore, we evaluated all mutations with a state-of-the-art deep learning-based scoring function called DDMut-PPI^13^ (figure 1C). This model is based on a Siamese network architecture with graph-based signatures and encodes both evolutionary and physical information. Each plot in figure 1 depicts the experimentally measured ΔΔ*G_Bindin_*_g_values, which are correlated against the computationally predicted values for each method. Our approach achieved a higher performance than FoldX in terms of Pearson correlation (0.59 compared to 0.47) and Root Mean Squared Error (RMSE) (1.69 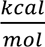 versus 1.98 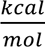). Notably, both methods produce large outliers in several cases that deviate strongly from the experimental data. For FoldX, some of these outliers are due to steric clashes, as the approach does not include energy minimization and solely relies on rotamer repacking. For both methods, several outliers are also related to mutations of charged residues and indicate poor overall modelling or wrong protonation state assignment (see also figure 2). Both FoldX and our method seem to perform on average better for mutations that lead to a lower binding affinity, hinting at an intrinsic bias from the crystal structures. In fact, many of the negative ΔΔ*G_Bindin_*_g_values in the SKEMPI v2.0 database are associated with the crystallized mutated variants. The binding affinity differences of these variants are typically the opposites of the positive values for the wildtype variants. These results might therefore be related to larger differences in sidechain packing or backbone conformation in the corresponding crystal structures. In contrast, DDMut-PPI shows a very good predictive performance on the SKEMPI v2.0 dataset, both in terms of Pearson correlation (0.92) and RMSE (0.82 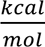). This result might be expected as the same dataset was also used to train and evaluate the DDMut-PPI method. Deep learning-based methods typically contain significantly more parameters than linear regression models and can therefore adapt very well to their training data. However, this carries the risk of overfitting to the training data and losing generalizability on unseen datasets as we show later. It remains to be determined what maximum level of predictive performance can be achieved by a generalizable deep learning-based scoring method on the SKEMPI v2.0 dataset, but related work suggests that these types of models require substantially larger datasets to be trained reliably^19^.

**Figure 1:**
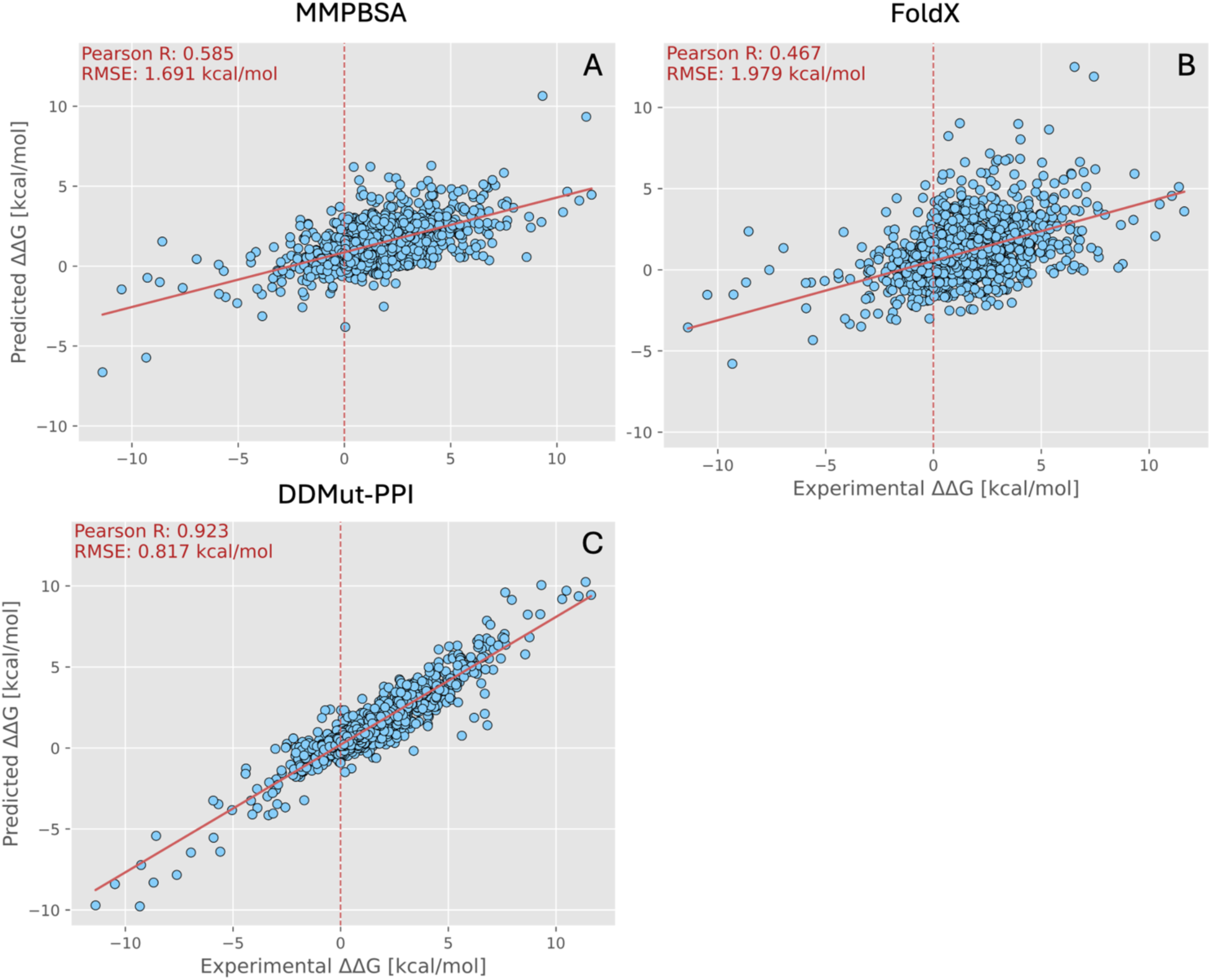
Correlations between predicted and experimentally determined binding free energy changes on the SKEMPI v2.0 dataset. Correlations are shown for three different methods: the MMPBSA-based scoring method presented in this work (A), FoldX (B) and DDMut-PPI (C).

**Figure 2:**
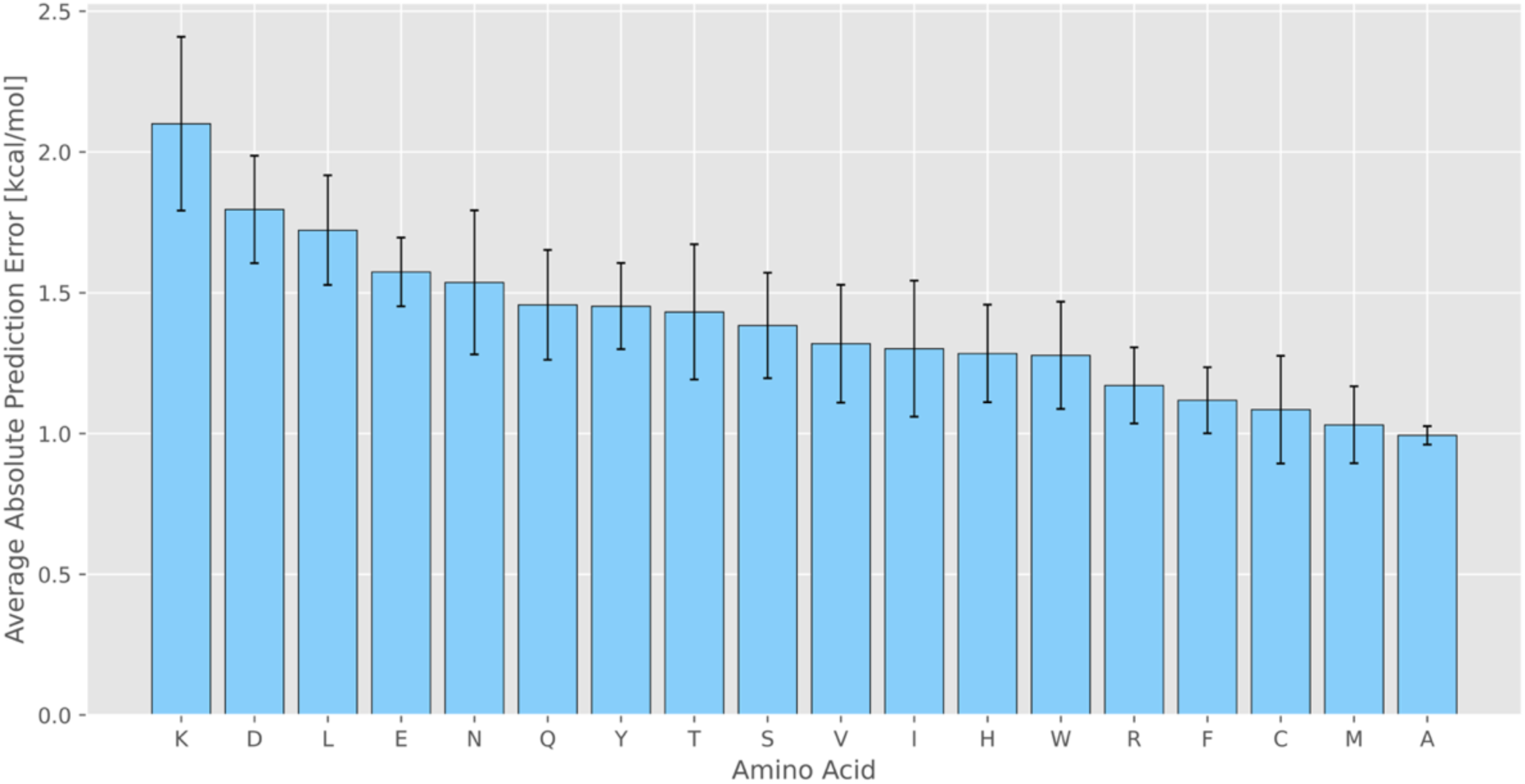
Average absolute binding free energy change prediction error of 18 different amino acid mutations (all except glycine and proline) on the SKEMPI v2.0 dataset for the MMPBSA-based scoring function presented in this work.

To identify potential biases in our method, we analyzed the average prediction error per type of amino acid mutation on the SKEMPI v2.0 dataset (see figure 2). The average prediction errors varied between about 1 to 2 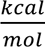. The largest average prediction error was observed for lysine mutations ( 2.1 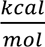), whereas arginine mutations exhibited substantially lower errors (1.2 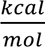) despite their comparable positive charge and side-chain flexibility. This suggests that the elevated prediction error for lysine mutations may primarily reflect characteristics of the underlying dataset, as these mutations are associated with larger average ΔΔ*G_Bindin_*_g_ values. Overall, large absolute ΔΔ*G_Bindin_*_g_ values tended to be associated with larger absolute errors.

Elevated errors were also observed for leucine mutations (1.7 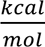) in comparison to isoleucine (1.3 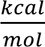) as well as for the negatively charged aspartic and glutamic acid (1.8 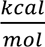 and 1.6 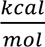), whereas alanine mutations showed only a small average prediction error of about 1.0 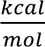 and a small variation in the average error. In addition, the hydroxyl-group containing amino acids tyrosine, threonine and serine as well as the amine-group containing amino acids asparagine and glutamine show errors of about 1.5 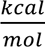. Together these results indicate that mutations involving charged and hydrogen-bonding sidechains remain particularly challenging.

The observed error pattern suggests two likely sources of uncertainty. First, inaccuracies in sidechain conformations generated during structural modeling (performed with FoldX, see methods) may affect the prediction of residue-specific interactions including charge-charge and hydrogen-bonding interactions. Second, the assignment of protonation states particularly for negatively charged amino acids (modelled as deprotonated, see methods) may contribute to errors in electrostatic interactions.

To evaluate model transferability beyond the training data, we compiled a dataset from a deep mutational scanning study of the SARS-CoV-2 spike receptor binding domain interacting with the human ACE2 receptor (see methods). This dataset provides both a stringent test to assess the generalizability of all methods and an opportunity to assess performance in a deep mutational scanning setting.

Figure 3 compares predicted and experimentally determined ΔΔ*G_Bindin_*_g_values for all three methods. The MMPBSA-based method achieved the best overall performance with a Pearson correlation coefficient of 0.51 and a RMSE of 0.97 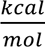 (figure 3A). In comparison FoldX yielded a lower correlation (0.40) and substantially higher RMSE ( 1.61 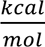) largely due to several pronounced outliers (figure 3B). The improved performance of the MMPBSA-based method suggests that the fitted energy terms capture mutation-induced binding affinity changes more accurately in the experimental system than the FoldX empirical energy function and shows that our model can generalize well outside of its training data.

**Figure 3:**
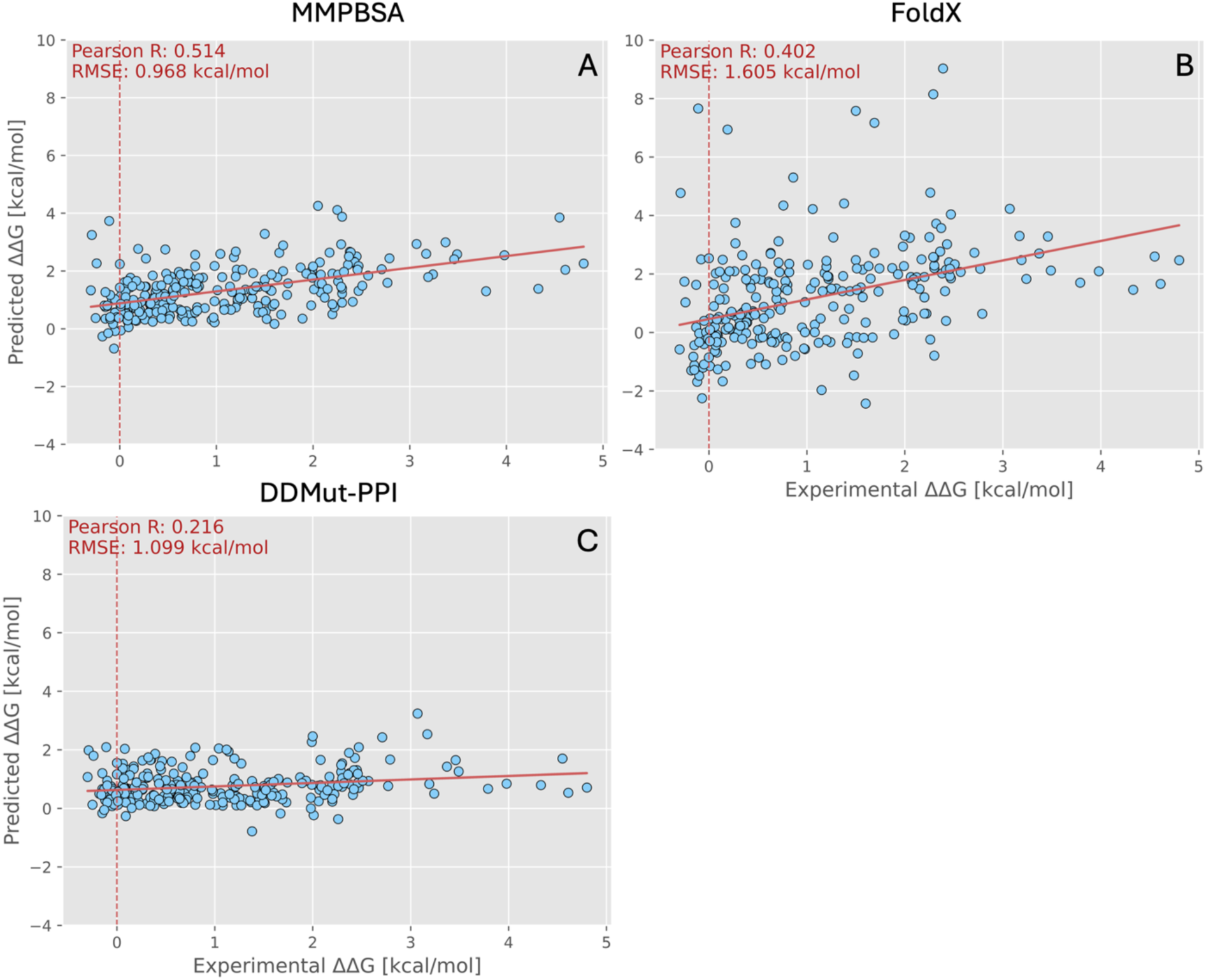
Correlations between predicted and experimentally determined binding free energy changes on the SARS-CoV-2 dataset. Correlations are shown for three different methods: the MMPBSA-based scoring method presented in this work (A), FoldX (B) and DDMut-PPI (C).

DDMut-PPI showed a very different performance (figure 3C). Although its RMSE (1.1 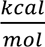) was a lot lower than the one of FoldX, the Pearson correlation coefficient was only 0.22 on the SARS-CoV-2 dataset, indicating limited ability to rank single residue variants according to their experimentally observed effects. Discrimination between favorable and adverse mutations is the primary goal in protein engineering and design, which is why correlation-based metrics are highly relevant in this context.

Notably, DDMut-PPI achieved a substantially higher performance on the SKEMPI v2.0 derived benchmark (Pearson correlation 0.92, see figure 1C), whereas both physics-based methods performed more consistently across the two datasets (see figure 1A and B).

As a third test case, we evaluated the methods on a binding affinity dataset of engineered dArmRPs with tailored specificity for different target peptides^34^. Previous computational studies of these protein-peptide complexes using three conceptually different physics-based protein design methods showed method-dependent biases with individual scoring functions performing well for some targets while showing reduced accuracy for others^7^. Since experimental reference structures were not available for all complex structures, all calculations were performed on computationally predicted models. To minimize uncertainties associated with model quality of the predicted structures, only the highest confidence structure was selected for the binding free energy calculations of each complex (see methods).

Figure 4 summarizes the evaluation of the dArmRP dataset using all three methods. In contrast to the SARS-CoV-2 benchmark, all three methods yielded relatively comparable predictive performance on this dataset, showing Pearson correlation coefficients of about 0.6 and above. Our method and FoldX (figure 4A and B) achieved slightly higher correlations of about 0.65. Interestingly, DDMut-PPI performs much better on the dArmRP dataset in terms of Pearson correlation than on the SARS-CoV-2 dataset (0.6 compared to 0.22, figure 4C). Notably, despite the similar correlation of our method and FoldX on this dataset, MMPBSA produces substantially fewer large outliers than FoldX, reflected in a significantly lower RMSE (0.73 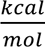 versus 1.56 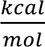). These results indicate that the MMPBSA-based scoring method provides more accurate affinity estimates while keeping comparable ranking performance.

**Figure 4:**
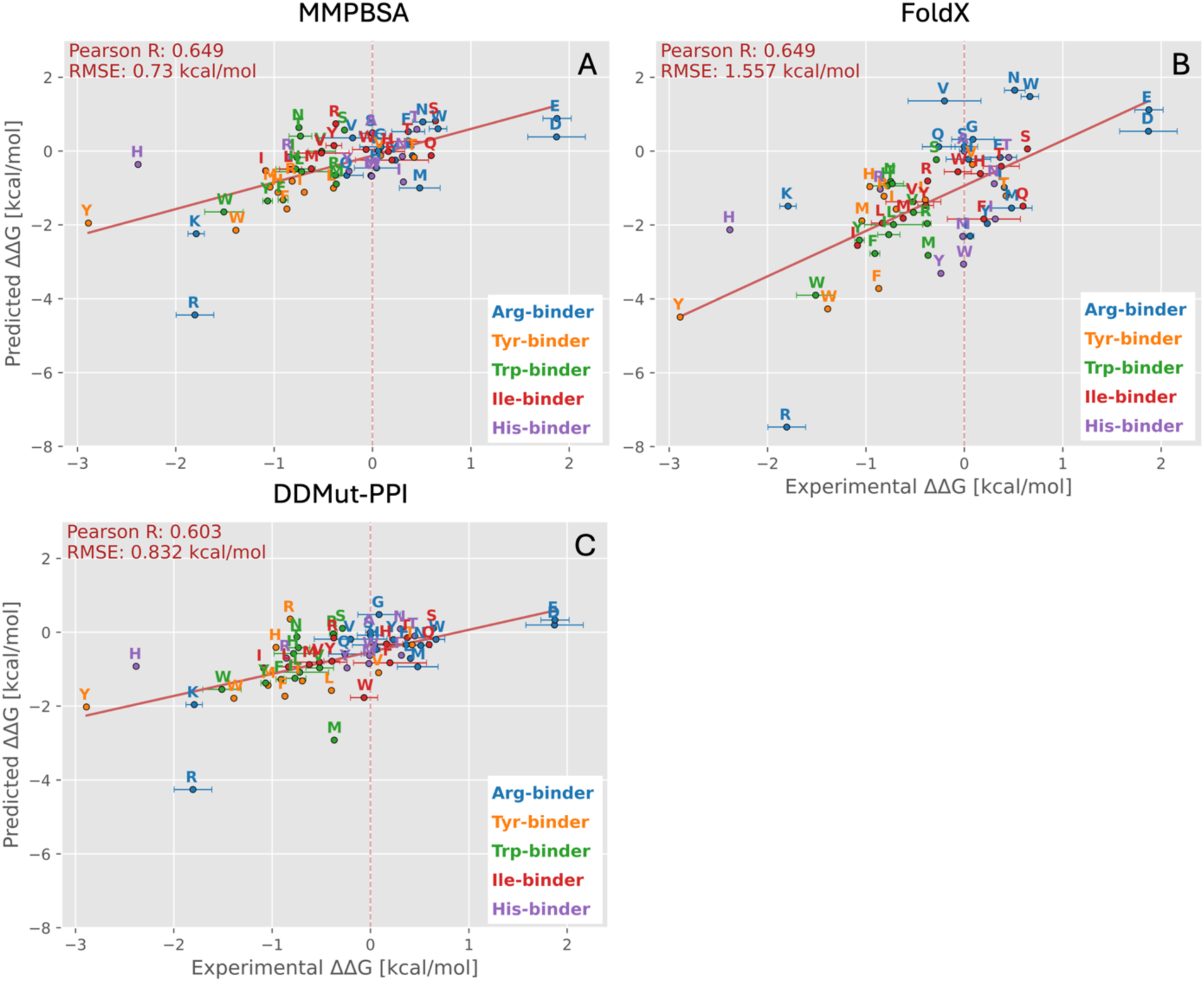
Correlations between predicted and experimentally determined binding free energy changes on the dArmRP dataset. Correlations are shown for the MMPBSA-based scoring method presented in this work (A), FoldX (B) and DDMut-PPI (C). Each plot shows five different dArmRPs, which are represented in varying colors. Each dArmRP was measured with a set of different peptides that differ by a single amino acid mutation, which is indicated in the plot.

Several trends were observed across all methods. In the Arg-binder, arginine affinities were consistently overpredicted relative to lysine, even though both residues show experimentally comparable binding affinities. Because the structural model was generated using an arginine-containing peptide, this tendency might reflect a bias that originates from the starting structure rather than the scoring functions themselves. However, this effect was most pronounced for FoldX. Another notable trend was observed for histidine binding to the His-binder, which was significantly underpredicted by both the MMPBSA-based method and DDMut-PPI. Inspection of the predicted protein-peptide complex revealed that the histidine sidechain was placed in an adjacent binding pocket rather than the experimentally expected site. This structural error could not be resolved during subsequent remodeling and therefore likely reflects limitations of the underlying structural model rather than deficiencies of the scoring functions. Overall, the results demonstrate that the MMPBSA-derived scoring function remains robust when applied to computationally predicted structures and achieves performance comparable to or better than alternative methods evaluated here.

### Conclusion

In this work, we developed a MMPBSA-based scoring function to predict the effects of point mutation on binding affinity at protein-protein and protein-peptide interfaces and compared it against the empirical force field FoldX and the recently developed deep learning-based method DDMut-PPI. While all three methods performed reasonably well on the training-related SKEMPI dataset, our MMPBSA-based approach showed the most consistent performance across the independent test sets. In particular, it achieved lower prediction errors than FoldX and maintained higher predictability than DDMut-PPI on the SARS-CoV2 benchmark. This highlights the importance of evaluating scoring functions beyond training-related datasets. Similar limitations in transferability of machine learning-based scoring functions have been reported previously^18^, suggesting that generalizability remains an important challenge for data-driven models. In contrast, our physics-based scoring function enables robust predictions of single mutation effects across diverse targets and for both experimentally determined and computationally predicted structures. Importantly, all evaluations presented here were based on individual minimized structures and do not rely on ensembles generated by molecular dynamics (MD) simulations. Although structural ensembles can improve the overall accuracy of empirical force fields for single mutation predictions^35,36^, these gains are often modest relative to the associated computational cost. The use of minimized structures therefore offers an attractive compromise between accuracy and efficiency. In return this typically requires high structural similarity between wildtype and mutant models to pinpoint mutational effects on the mutation sites. Among other factors, differences in structural comparability may explain why MMPBSA calculations based on minimized structures previously showed higher correlations with experiment than those based on ensembles generated from short MD simulations^11,37–39^.

Overall, our results demonstrate that MMPBSA-derived energy terms provide a robust basis for predicting mutation-induced binding affinity changes. The new framework introduces a generalizable physics-based alternative to existing empirical and machine-learning approaches and may prove useful for protein engineering, interface optimization and the assessment of mutation effects in diverse protein complexes.

## Materials and methods

### Dataset compilation

Three datasets were analyzed in this work. The first dataset was compiled from the SKEMPI v2.0 database^31^ and consists of 1662 datapoints that belong to 175 wildtype crystal structures. To compile the final dataset, all duplicate data points were first merged by averaging their binding affinity values. This refers to datapoints originating from different studies or measured under different conditions. Second, all mutations were removed that are not located at the protein-protein binding interface (defined as core, rim, support region according to Levy^40^). Third, all data points corresponding to structures other than crystal structures, as well as those for which the crystal structure had a resolution worse than 2.5 Å, were removed. Fourth, all datapoints were removed that involve glycine or proline mutations, as these mutations can lead to large secondary structure or entropy changes, which cannot be properly reflected in our modelling and scoring approach. The second dataset was compiled from an exhaustive mutagenesis study on the RBD of the SARS-CoV-2 spike glycoprotein in interaction with the human ACE2 receptor and consists of 272 datapoints belonging to a single wildtype crystal structure^32^. From the original dataset, which consists of 3684 datapoints, all mutations involving glycine or proline were removed and only mutations at the binding interface were retained (following the same definition as above). In line with our previous work the third dataset consists of 63 measured binding affinities for a set of five dArmRP-binders together with a set of diverse peptides^7^. ΔΔ*G_Bindin_*_g_ values were calculated using either an alanine reference for the dArmRP dataset, or the wildtype structure for the set of SKEMPI/SARS-CoV-2 mutants. When available, the errors of the experimental values are shown, typically representing the fitting error of the measured binding curve. The ΔΔ*G_Bindin_*_g_ values upon mutation were in all cases defined as:

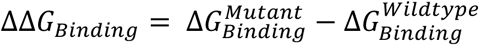

### Structural modelling

Wildtype crystal structures were selected as provided by the SKEMPI database, the crystal structure with the PDB-ID: 6M0J (Resolution: 2.45 Å)^41^ was used as a wildtype structure for all modelling related to the SARS-CoV-2 dataset. Crystal structures were prepared by first removing all waters, crystallizing agents and other non-protein molecules. If there were protein chains that are not involved in the binding, these were also removed. For the dArmRP-peptide complexes, consensus sequences for protein and peptide were used as a basis for modelling. These sequences were taken from Stark et al.^34^. The last four residues of the protein and the last two residues of the peptide were removed, as they were highly flexible in the structure predictions and thus lowered the overall quality of the predictions. However, these residues are far away from the binding pocket of interest and are thus not expected to have a significant impact on the results. The respective armadillo pocket residues were exchanged at positions: 190, 194, 197, 229, 232, 233 and 236 of the protein and the respective peptide residue was exchanged at position 4 of the peptide. Each Armadillo-peptide complex was modelled with the amino acid that shows experimentally the highest binding affinity in the respective binding pocket of interest^34^. Modelling of the complexes was carried out using the Chai-1 structure prediction method^42^. Chai-1 was run with 3 trunk recycles and 200 diffusion steps, Evolutionary Scale Modeling (ESM) language model embeddings^43^ as well as Multiple Sequence Alignments (MSAs) and template structures were utilized in all structure predictions. Five diffusion samples were generated, of which the best according to the highest Chai-1 aggregate score was selected for further modelling.

The FoldX command *RepairPDB* was first run to repack the structures and optimize their energy inside the FoldX 5.1 force field^10^. Mutated protein models were then built based on the repacked structures by running the command *BuildModel*. It should be noted that FoldX generates a corresponding wildtype model for each mutated model, in which the same residues are repacked to ensure structural consistency. Accordingly, for each mutant the corresponding wildtype model generated by FoldX was used as a reference structure. Hydrogen atoms were then added and their positions optimized using *reduce*^44^. All carboxylic groups were modelled as deprotonated and all amine groups as fully protonated. For histidine residues protonation states were determined based on optimal hydrogen-bonding interactions. Wildtype and mutant models were then minimized with a harmonic restraining force of 10 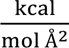 applied to all heavy atoms of the protein backbone. Minimization was carried out until energy convergence. The AMBER ff19SB force field^45^ and OBC2 implicit solvent model^46^, as implemented in OpenMM^47^, were employed in all minimizations. Hydrogen bond lengths were restrained using the SHAKE algorithm^48^.

### Binding free energy calculations

The change in free energy upon binding was calculated in all cases by subtracting the free energy of the receptor and ligand from the free energy of the complex. Here, receptor and ligand refer either to a single or a group of polypeptide chains, each defining the two binding partners A and B. The conformations for the unbound states (receptor and ligand) were assumed to be the same as in the bound state (complex) and no additional conformations were generated.

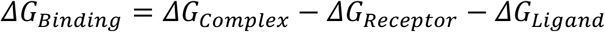

The individual binding free contributions that were calculated are: the VdW energy, the Coulomb energy, the polar and the nonpolar solvation energy. We additionally included the folding stability change, which was calculated by running the FoldX command *Stability*. We decided to combine the Coulomb and the polar solvation energy into a single electrostatic energy term in our scoring function. The nonpolar solvation energy was estimated based on the change in buried solvent accessible surface area (SASA) upon binding. The VdW and coulomb energy were calculated based on the Lennard-Jones 12-6 potential and based on Coulomb’s Law assuming a protein dielectric constant of 4.0. AMBER parameters were used for all atoms, and the calculations were performed using OpenMM. To calculate the free energy change of solvation the linearized Poisson-Boltzmann equation was solved using the Adaptive Poisson-Boltzmann Solver (APBS)^49^. In all calculations the protein dielectric constant was set to 4.0, and the solvent dielectric constant was set to 80.0. The salt concentration was set to 150 mM monovalent salt with an ion exclusion radius of 2.0 Å. The number of grid points in each dimension was determined to be 10 Å larger than the size of the solute. The grid resolution was set to 0.5 Å and the charges of the solute were mapped to the grid using a cubic B-spline discretization. The solvent probe radius was set to 1.4 Å and the dielectric boundary was defined as the solvent excluded surface. A “vacuum state” without ions and with an external dielectric of 4.0 was used as a reference state and subtracted from the solvated state to obtain the solvation free energy change. AMBER partial atomic charges and mbondi2 radii^46^ were used for all atoms.

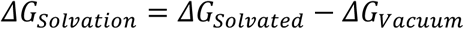

For FoldX-based binding free energy calculations, the command *AnalyseComplex* was run, which calculates the free energy of binding in a similar way by unfolding the unbound binding partners A and B from the bound state and subtracting their energies from that state (see above). All FoldX energy calculations are based on the FoldX 5.1 force field^10^. For all evaluations based on DDMut-PPI^13^, the webserver available under the URL: https://biosig.lab.uq.edu.au/ddmut_ppi was used. The same PDB files together with the mutations were submitted to the webserver and the results obtained.

### Model evaluation

We used the Pearson correlation coefficient to assess the linear relationship between the computationally predicted and experimentally determined ΔΔ*G_Bindin_*_g_ values. The calculation was performed based on the following equation:

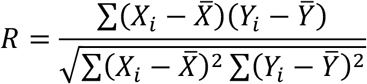

In the equation *X_i_* and *Y_i_* represent individual datapoints in both datasets *X* and *Y* and *X̅* and *Y̅* represent the mean values of both datasets *X* and *Y*. We further accessed the average error of the predictions using the RMSE:

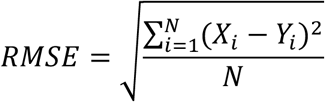

Here, *X_i_* and *Y_i_* represent individual datapoints in both datasets *X* and *Y* and *N* represents the total number of datapoints in both datasets.

## Acknowledgements

This work was supported by the European Innovation Council EIC Transition grant 10105802 “PRe-ART-2T”. Thibaud Lepoivre was supported through the Erasmus+ programme of the European Union. We thank members of the Höcker lab and the PReART research team for discussions.

